# Optimizing anesthesia and delivery approaches for dosing into lungs of mice

**DOI:** 10.1101/2023.02.01.526706

**Authors:** Yurim Seo, Longhui Qiu, Mélia Magnen, Catharina Conrad, S. Farshid Moussavi-Harami, Mark R Looney, Simon J Cleary

## Abstract

Microbes, toxins, therapeutics and cells are often instilled into lungs of mice to model diseases and test experimental interventions. Consistent pulmonary delivery is critical for experimental power and reproducibility, but we observed variation in outcomes between handlers using different anesthetic approaches for intranasal dosing into mice. We therefore used a radiotracer to quantify lung delivery after intranasal dosing under inhalational (isoflurane) versus injectable (ketamine/xylazine) anesthesia in C57BL/6 mice. We found that ketamine/xylazine anesthesia resulted in delivery of a greater proportion (52±9%) of an intranasal dose to lungs relative to isoflurane anesthesia (30±15%). This difference in pulmonary dose delivery altered key outcomes in models of viral and bacterial pneumonia, with mice anesthetized with ketamine/xylazine for intranasal infection with influenza A virus or *Pseudomonas aeruginosa* developing more robust lung inflammation responses relative to control animals randomized to isoflurane anesthesia. Pulmonary dosing efficiency through oropharyngeal aspiration was not affected by anesthetic method and resulted in delivery of 63±8% of dose to lungs, and a non-surgical intratracheal dosing approach further increased lung delivery to 92±6% of dose. Use of either of these more precise dosing methods yielded greater experimental power in the bacterial pneumonia model relative to intranasal infection. Both anesthetic approach and dosing route can impact pulmonary dosing efficiency. These factors affect experimental power and so should be considered when planning and reporting studies involving delivery of fluids to lungs of mice.

**Graphical abstract:** 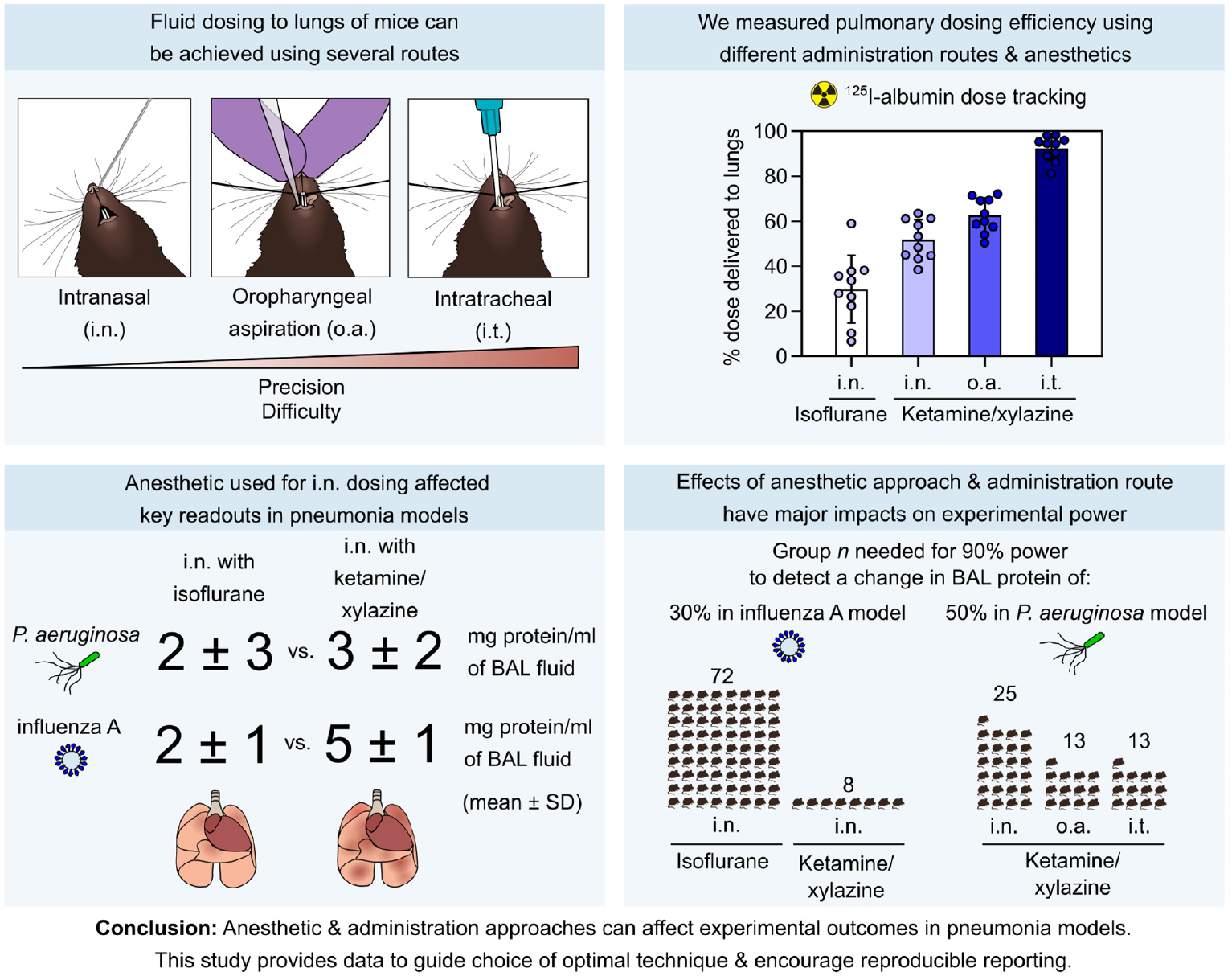

## Introduction

Studies investigating lung infections, lung injury, allergic airway inflammation, lung fibrosis, lung cancer, and lung stem cell biology often require delivery of experimental agents to lungs of mice. Administration routes for bolus dosing of fluids into lungs include intranasal (i.n.) dosing, intratracheal (i.t.) dosing and dosing through oropharyngeal aspiration (o.a.). Choice of dosing route is an important decision in study design as experimental outcomes can be altered by the quantity of dose delivered to lungs or to extrapulmonary tissues. Different dosing routes also vary in anesthetic requirements, invasiveness, and technical difficulty.

To guide experimental approach, a previous study assessed the effect of various factors including type of anesthetic on the distribution of i.n. doses into BALB/c mice. This study concluded that either injectable (Avertin) or inhaled (isoflurane, halothane) anesthetics resulted in similar delivery to lungs (1). Since this influential report, several factors have changed. Safety concerns have led to a decline in use of both Avertin and halothane (2, 3). Increased availability of knockouts and transgenics on the C57BL/6 (B6) background has led to B6 mice becoming the most widely used laboratory strain. Additionally, minimally invasive approaches for dosing via o.a. and i.t. routes have been developed which can more efficiently deliver fluids to lungs relative to i.n. dosing (4–6).

In our previous work we noticed that i.n. doses passed more readily into the nostrils in studies where B6 background mice were anesthetized with ketamine/xylazine compared to experiments in which mice were anesthetized with isoflurane (7, 8), but we did not know whether anesthetic used during dosing was affecting pulmonary deposition. To guide future studies using these mice and anesthetics, we therefore measured the effect of anesthetic approach on i.n. delivery of fluid to lungs of B6 mice. As we have used and refined o.a. and i.t. methods, we also measured dose distribution using these administration routes.

We found striking effects of both anesthetic approach and dosing route on the efficiency of pulmonary delivery. Our results will be useful to guide design of experiments with improved reproducibility – an international biomedical research policy goal (9, 10). Additionally, our findings indicate potential strategies to reduce the number of mice needed to produce clear results from experiments involving dosing of fluids into lungs.

## Methods

### Animals

C57BL/6 background mice (Jax #000664) were housed at the UCSF Parnassus Laboratory Animal Resource Center specific pathogen-free facility. Male and female mice were used in equal numbers per group at ages 6-14 weeks. Mice were kept on a 12-hour light-dark cycle. Protocols were approved by the UCSF Institutional Animal Care and Usage Committee.

### Anesthesia

Mice were anesthetized either by inhalation of isoflurane (4% in oxygen) or by intraperitoneal (i.p.) injection with ketamine (70 mg/kg) and xylazine (15 mg/kg) in normal saline. For fluid dosing to lungs, it is important not to overdose ketamine/xylazine as higher doses can cause asphyxiation from aspirated fluid. For terminal anesthesia prior to collection of lung samples, mice were euthanized with ketamine (100 mg/kg) and xylazine (40 mg/kg) prior to exsanguination.

### Intranasal dosing

Gel-loading pipette tips (Sorenson #13810) were used to introduce 50 μl of dose dropwise into the posterior opening of one nare. Mice were held upright for 20 seconds after dosing to allow aspiration of dose.

### Tracking radiolabeled albumin doses

For quantitative and visual tracking of inoculum we used 50 μl of phosphate-buffered saline (PBS) containing ^125^I-albumin (0.25 mg/ml, ∼2.5 KBq/ml, Jeanatope, Iso-Tex Diagnostics, Inc.) and Evans blue dye (1 mg/ml). Organ samples were collected 10 minutes after mice were dosed. Dose distribution was measured using a gamma counter (Packard 5000 series) against three standards containing 100% of injected dose.

### Influenza A virus infection model

Mice were infected by the i.n. route with 50 plaque-forming units (p.f.u.) of influenza A virus (A/PR/8/34 H1N1) propagated with Madin-Darby canine kidney (MDCK) cells. For propagation, MDCK cells cultured in minimal essential medium (MEM) supplemented with 10% FBS and penicillin/streptomycin in a humidified incubator at 37ºC and 5% CO_2_ were infected and cultured for 72h. The supernatant containing the virus was then collected and stored at -80ºC. Infectious virus was quantified by culturing dilutions of the viral stock with MDCK cells in a 6-well plate for 1 hour, followed by addition of an overlay of 1.2% Avicel RC-581 in MEM, culture for 72 hours, formalin fixation and staining with crystal violet for p.f.u. determination (11). Viral stocks were diluted in sterile PBS at 4ºC prior to inoculation. Mice were dosed at zeitgeber time (ZT) 3-5 and handled under biosafety level 2 conditions.

### Bronchoalveolar lavage analysis

After terminal anesthesia and exsanguination, lungs were collapsed by opening the diaphragm and tracheal insertion of 20G stub needles. A 1 ml syringe containing 1 ml of phosphate-buffered saline was then washed in and out of the lungs three times to recover bronchoalveolar lavage (BAL) fluid. BAL cells were counted using a LUNA-II automated cell counter (Logos Biosystems) and BAL supernatant total protein was measured using a Pierce total protein assay (Thermo Scientific, #23225).

### Dosing by oropharyngeal aspiration

As previously described, anesthetized mice were placed on an intubation platform suspended by their upper incisors with the tongue gently pulled out of the mouth (5, 12). The fluid dose was then pipetted directly onto the distal oropharynx at 50 μl volume with both nares covered to obligate breathing through the mouth. After ∼30 seconds, mice were removed from the platform and placed supine until sample collection.

### Non-surgical intratracheal dosing

Customized intubation and injection apparatus was prepared from a blunted 22G 1” Safelet IV Catheter (Nipro, #CI+22225-2C) customized into an endotracheal tube, and a blunted 28G insulin syringe (BD #329461) attached to PE-10 tubing (**Figure 4A**). Cushions were added using cyanoacrylate glue and PU-40 or PE-20 tubing. Mice were positioned as with o.a. dosing, with transillumination and adjustment of body position used for visualization of the larynx (**Figure 4B**). Mice were then orotracheally intubated with correct placement confirmed by attaching a manometer to the endotracheal tube and checking for oscillation of water column with breathing movements (**Figure 4C**) (13). The tubing attached to the syringe was then inserted into the catheter (as shown in **Figure 4D**) for injection of dose at 50 μl volume followed by 120 μl air.

### Pseudomonas aeruginosa infection model

*Pseudomonas aeruginosa* (PAO1, ATCC #BAA-47) was grown to logarithmic phase in suspension in tryptic soy broth (TSB). Pellets of PAO1 were then resuspended in PBS at 4°C and adjusted to a density of 1×10^6^ colony forming units (c.f.u.) per 50 μl inoculation volume for mouse infections.

Infected mice were given a subcutaneous dose of 0.5 ml of normal saline at 4 hours after infection as fluid support. BAL fluid was collected as described above at 24 hours after infection.

### Experimental design and statistical analysis

Mice were randomly assigned to groups with blocking by cage, and samples were collected and quantified with investigators blinded to groups. For infection studies, the handler dosing mice was also blinded during dosing, with a second unblinded handler in control of anesthesia. Group *n* was set prior to study initiation and analysis. Where necessary, data were transformed prior to statistical testing according to distribution. Statistical analyses used InVivoStat 4.4 (body weight and power analysis) or GraphPad Prism 9 (other comparisons). The tests used for each analysis are stated in figure legends with *P*=0.05 as α threshold. Data are reported as means ± standard deviation unless otherwise stated.

## Results

In previous experiments we noticed that B6 mice anesthetized with ketamine/xylazine smoothly aspirated i.n. doses, whereas i.n. doses sometimes bubbled back out of the nares of isoflurane-anesthetized mice (7, 8). As previous studies assessing effects of anesthesia on i.n. delivery to lungs used anesthetics or mouse strains not used in our protocols (1, 14, 15), we aimed to determine whether use of isoflurane or ketamine/xylazine anesthesia during i.n. dosing affected delivery of dose to lungs of B6 mice.

We found that relative to isoflurane anesthesia, use of ketamine/xylazine anesthesia during i.n. dosing resulted in delivery of dose to more distal regions of lung (**Figure 1A**) and increased pulmonary dosing efficiency (**Figure 1B**). The proportion of dose not delivered to the lungs was not immediately swallowed, but either remained in the upper respiratory tract or was refluxed out of the nostrils (quantified as **“**Other” in **Figure 1B**).

**Figure 1.**
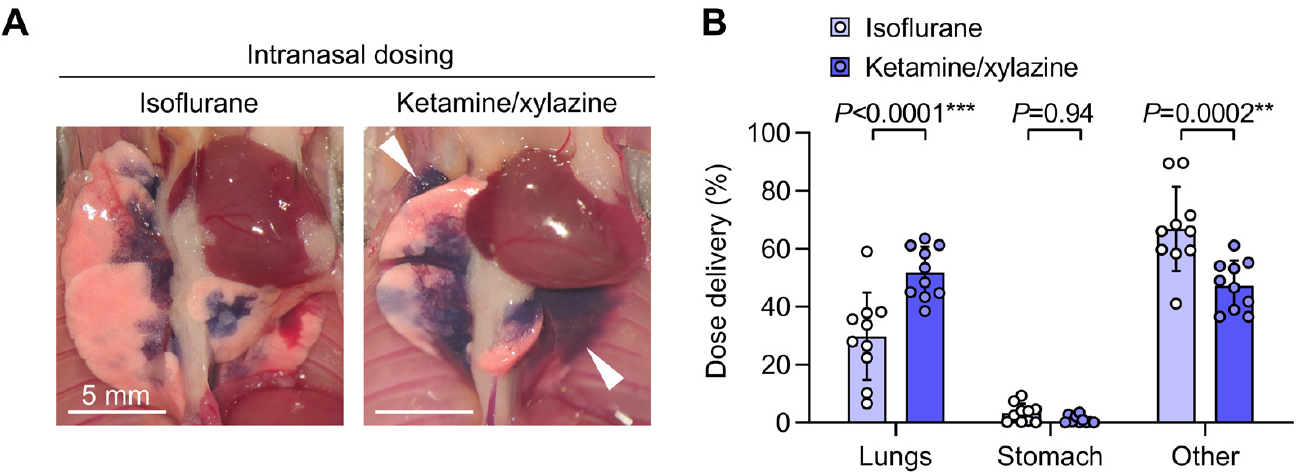
Increased pulmonary dosing efficiency with ketamine/xylazine versus isoflurane anesthesia during intranasal dosing. A. B6 mice were given intranasal (i.n.) doses containing ^125^I-albumin radiotracer and Evans blue dye under either isoflurane or ketamine/xylazine anesthesia. Photographs show lungs after euthanasia and thoracotomy, with delivery of dye to more distal regions of the lungs with ketamine/xylazine anesthesia (white arrowheads). B. Effect of anesthetic type on dose distribution quantified using radiotracer. Means ± standard deviation, n=10. *P*-values are from a repeated measures two-way ANOVA with Holm-ŠÍdák tests for effect of dosing route within each location.

The i.n. dosing route is in widespread use in respiratory virus infection models. Current protocols suggest that handlers can use either isoflurane or ketamine/xylazine anesthesia during i.n. infection with influenza A virus (16). We therefore formally tested whether the effect of anesthetic approach on intranasal dosing to lungs could be a factor altering outcomes and reproducibility of studies of respiratory viral infection.

With one handler delivering anesthesia, and a second handler blinded to anesthetic approach dosing and assessing mice, we gave B6 mice randomized to isoflurane or ketamine/xylazine anesthesia prior to i.n. doses containing 50 p.f.u. of PR8 influenza A virus.

We observed bubbling of dose back out of nares and down the philtrum in mice in our biodistribution study. During infection with PR8, the handler blinded to anesthesia approach therefore recorded whether dose reflux was observed. We found that isoflurane-anesthetized mice consistently refluxed some dose back out of their nares, whereas mice anesthetized with ketamine/xylazine smoothly aspirated doses without visible reflux (**Figure 2A,B**).

**Figure 2.**
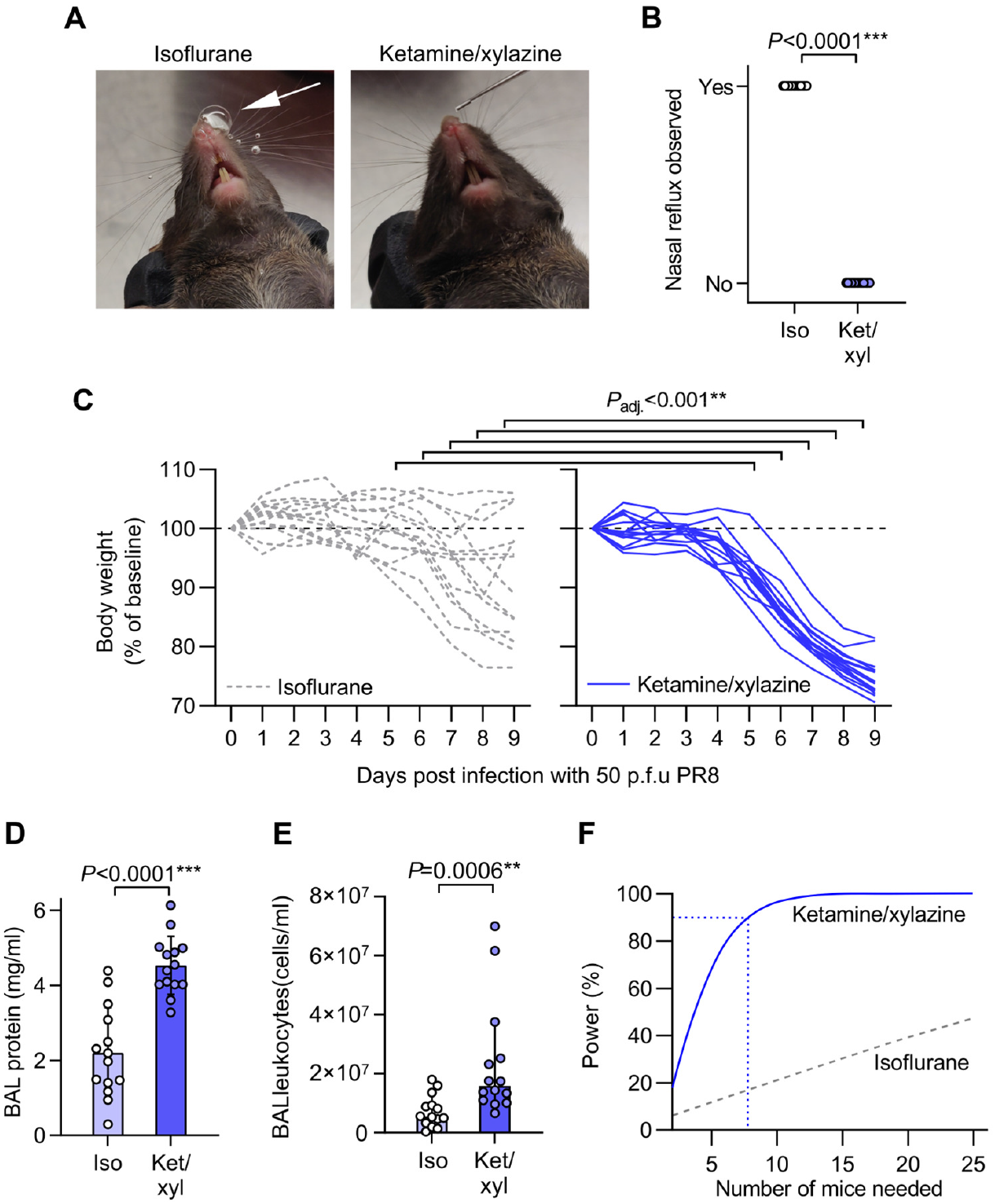
Increased body weight loss and lung inflammation after intranasal infection with influenza A virus under ketamine/xylazine relative to isoflurane anesthesia. **A**. Mice were randomized to receive either isoflurane or ketamine/xylazine anesthesia for intranasal dosing with 50 p.f.u. of H1N1 influenza A virus A/PR/8/1934 (PR8). Photographs of mice show presence of nasal reflux during intranasal dosing under isoflurane anesthesia (white arrow) but not ketamine/xylazine anesthesia. B. Quantification of incidence of nasal reflux during intranasal dosing under isoflurane (iso) versus ketamine/xylazine (ket/xyl). C. Body weight changes over 9 days post infection. D. BAL supernatant protein concentration at day 9 post infection. E. Leukocyte counts from BAL fluid at day 9 post infection. F. Output of power analysis using total protein data in Figure 2D to estimate number of mice needed per group to detect a 30% change in BAL protein concentration using an unpaired two-tailed t-test. Means ± standard deviation except for **E** which shows medians ± 95% confidence intervals and was log_10_-transformed prior to analysis, n=14. *P*-values are from: **B**: Fisher’s exact test, **C**: repeated measures mixed model approach with baseline values as covariates and adjusted (adj.) *P*-values from Holm’s tests for effect of anesthesia within each time point; **D**,**E**: unpaired two-tailed t-tests.

We also monitored body weight daily as an index of general health status. All mice anesthetized with ketamine/xylazine at time of infection had lost weight at day 9, but weight loss was significantly lower in the isoflurane-anesthetized group from 5 to 9 days post infection, with some mice in the isoflurane group gaining weight after inoculation (**Figure 2C**).

At 9 days post infection we collected bronchoalveolar lavage (BAL) fluid from infected mice to measure vascular leak and leukocyte recruitment into lung airspaces as indices of lung inflammation. Both supernatant protein concentration and leukocyte counts were higher in BAL fluid from ketamine/xylazine-anesthetized mice compared to isoflurane-anesthetized mice (**Figure 2D,E**).

Using the BAL protein data in **Figure 2D** we ran a power analysis to determine the group size needed for future experiments aimed at detection of a 30% change in BAL fluid protein concentration using unpaired two-tailed t-tests. We found that 9 mice per group would be needed to run such an experiment with 95% power with ketamine/xylazine anesthesia (**Figure 2F**). In comparison, the isoflurane anesthesia approach would likely not be feasible for experimental use as an experiment with 25 mice per group would still have less than 50% power (**Figure 2F**).

Together, these results indicate that isoflurane anesthesia spares a reflex involving sensing of fluid in the upper airways and limitation of pulmonary aspiration. In contrast, use of ketamine/xylazine anesthesia circumvents this reflex, facilitating aspiration of a greater proportion of i.n. dose. This effect means that the two anesthesia approaches yield different efficiency and distribution of i.n. dosing efficiency to the lungs, affecting key outcomes in a respiratory virus infection model.

Compared with i.n. dosing, the o.a. route involving aspiration from the distal oropharynx can result in less exposure of nasal sinuses to inoculum and increased dosing efficiency to the lungs. Since anesthetic type affected i.n. dosing, we sought to also determine whether different anesthesia approaches altered delivery of o.a. doses to the lungs.

Breath-holding responses were observed in some isoflurane-anesthetized mice after doses were dropped onto the oropharynx, but isoflurane-anesthetized mice eventually aspirated doses with nasal reflux and swallowing prevented by covering the nares and retracting the tongue. Breath holding was not observed in ketamine/xylazine anesthetized mice given o.a. doses, potentially resulting in more rapid aspiration over multiple breaths and patchier deposition (**Figure 3A**). Tracking dose delivery quantitatively, we did not detect any effect of anesthesia approach on o.a. dose deposition in the lungs (**Figure 3B**).

**Figure 3.**
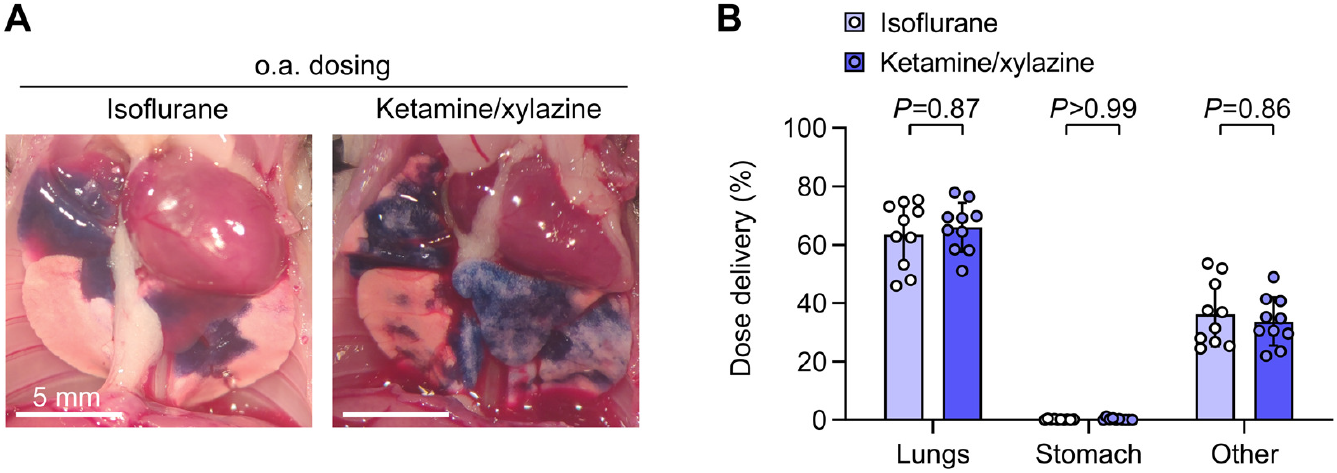
Pulmonary dosing efficiency using oropharyngeal aspiration is not altered by anesthetic type. A. B6 mice were given intranasal (i.n.) doses containing ^125^I-albumin radiotracer and Evans blue dye under either isoflurane or ketamine/xylazine anesthesia and euthanized 10 minutes later. Assessment of lungs after thoracotomy showed bilateral delivery of dye to distal regions of lungs with both anesthetic types. B. No effect of anesthesia approach was found on biodistribution of radiotracer. Means ± standard deviation, n=10. *P*-values are from a repeated measures two-way ANOVA with Holm-ŠÍdák tests for effect of dosing route within each location.

We conclude from this study that anesthetic approach is therefore unlikely to have a major impact on dosing to the lungs via the o.a. route.

Non-surgical i.t. dosing approaches have potential for more precise lung dosing relative to i.n. and o.a. dosing. Previous studies suggest that o.a. dosing can yield similar dosing efficiency compared to i.t. dosing, but these reports have not directly measured lung delivery using the latest non-surgical i.t. approaches (17–20). We have optimized an approach for i.t. dosing involving direct visualization of the larynx, orotracheal intubation with customized catheter, confirmation of airway placement using a manometer, and then injection using a customized syringe (5, 6, 12, 13) (**Figure 4A-D**), We therefore sought to measure pulmonary dosing efficiency using our non-surgical approach for i.t. dosing, comparing to a control group dosed with the o.a. approach, using ketamine/xylazine anesthesia.

**Figure 4.**
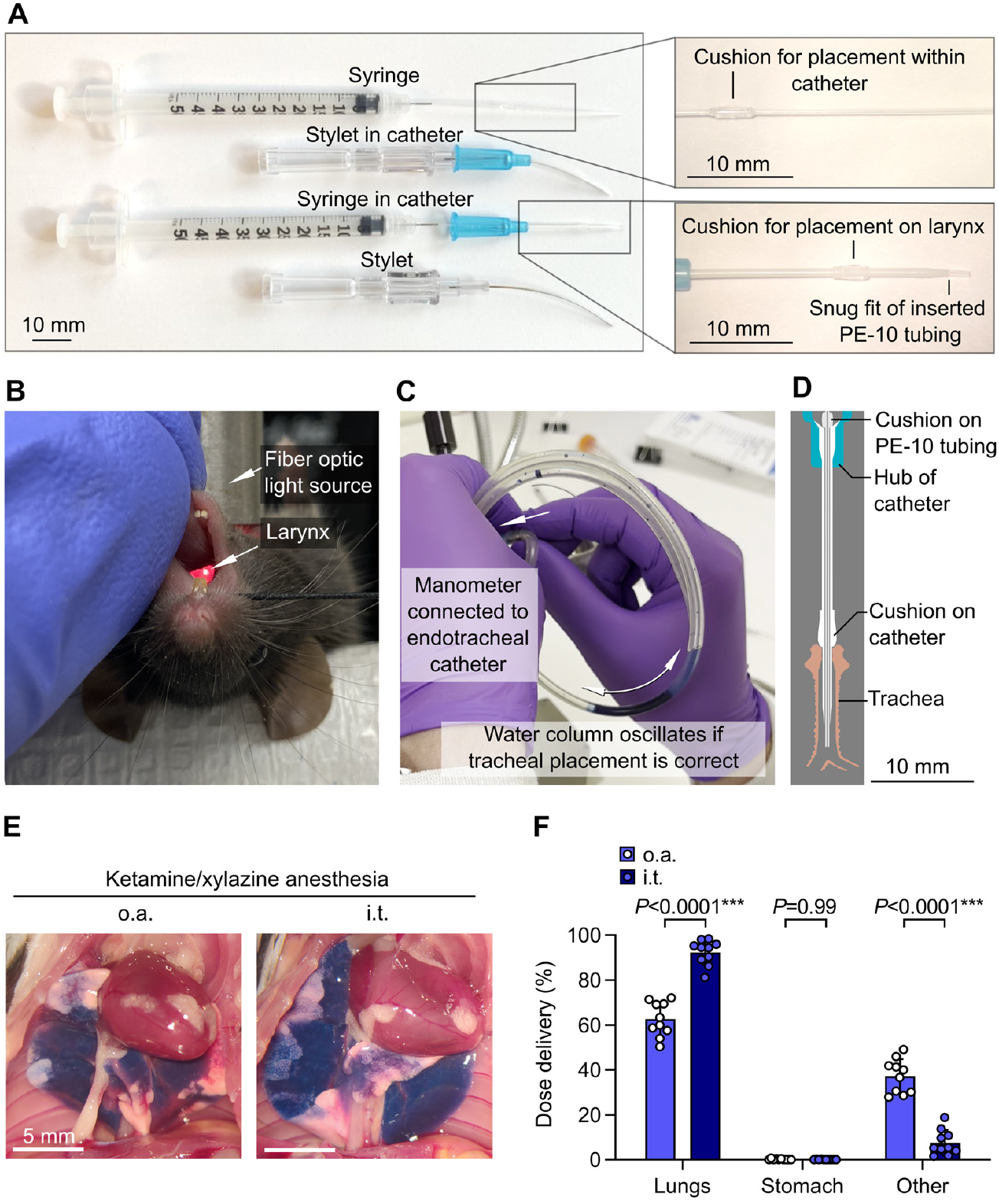
Increased pulmonary dosing efficiency with non-surgical intratracheal dosing relative to oropharyngeal aspiration. A. Customized catheters and syringes for orotracheal intubation and precise intratracheal injections through endotracheal tube. B. Mouse on intubation platform showing direct visualization of laryngeal inlet through transillumination the trachea. C. Manometer used for confirmation of correct airway placement after orotracheal intubation. D. Drawing showing cross section of tubing, catheter and trachea. D. B6 mice were anesthetized with ketamine/xylazine and then given ^125^I-albumin and Evans blue by either oropharyngeal aspiration (o.a.) or intratracheal (i.t.) routes. Photographs show representative dose distribution. F. Effect of administration approach on dose distribution quantified using radiotracer. Means ± standard deviation, n=10. *P*-values are from a repeated measures two-way ANOVA with Holm-ŠÍdák tests for effect of dosing route within each location.

We found that i.t. dosing yielded increased pulmonary dose deposition relative to o.a. dosing (**Figure 4E,F**). This result is indicative that although o.a. and i.t. routes deliver the majority of injected dose to the lungs, i.t. dosing might be desirable in situations where precise dosing to lungs is needed.

We have previously modeled bacterial lung infections by infecting mice by the i.n., o.a. and i.t. routes (21–24). To guide future studies using bacterial pneumonia models and directly compare performance of each of the dosing methods assessed above, we studied the effects of administration approach on lung inflammation following infection with *Pseudomonas aeruginosa*, a bacterium which causes ventilator-associated pneumonia and lung infections in cystic fibrosis.

Similar to our data from the influenza A infection model, i.n. dosing with the PAO1 isolate of *P. aeruginosa* under isoflurane anesthesia resulted in less influx of protein and cells into the bronchoalveolar space than with ketamine/xylazine anesthesia (**Figure 5A, B**). One mouse in the isoflurane group developed a very high BAL protein response, driving increased variance and an even greater difference in estimated power between the two anesthesia approaches than we calculated in the viral infection model (**Figure 5C**).

**Figure 5.**
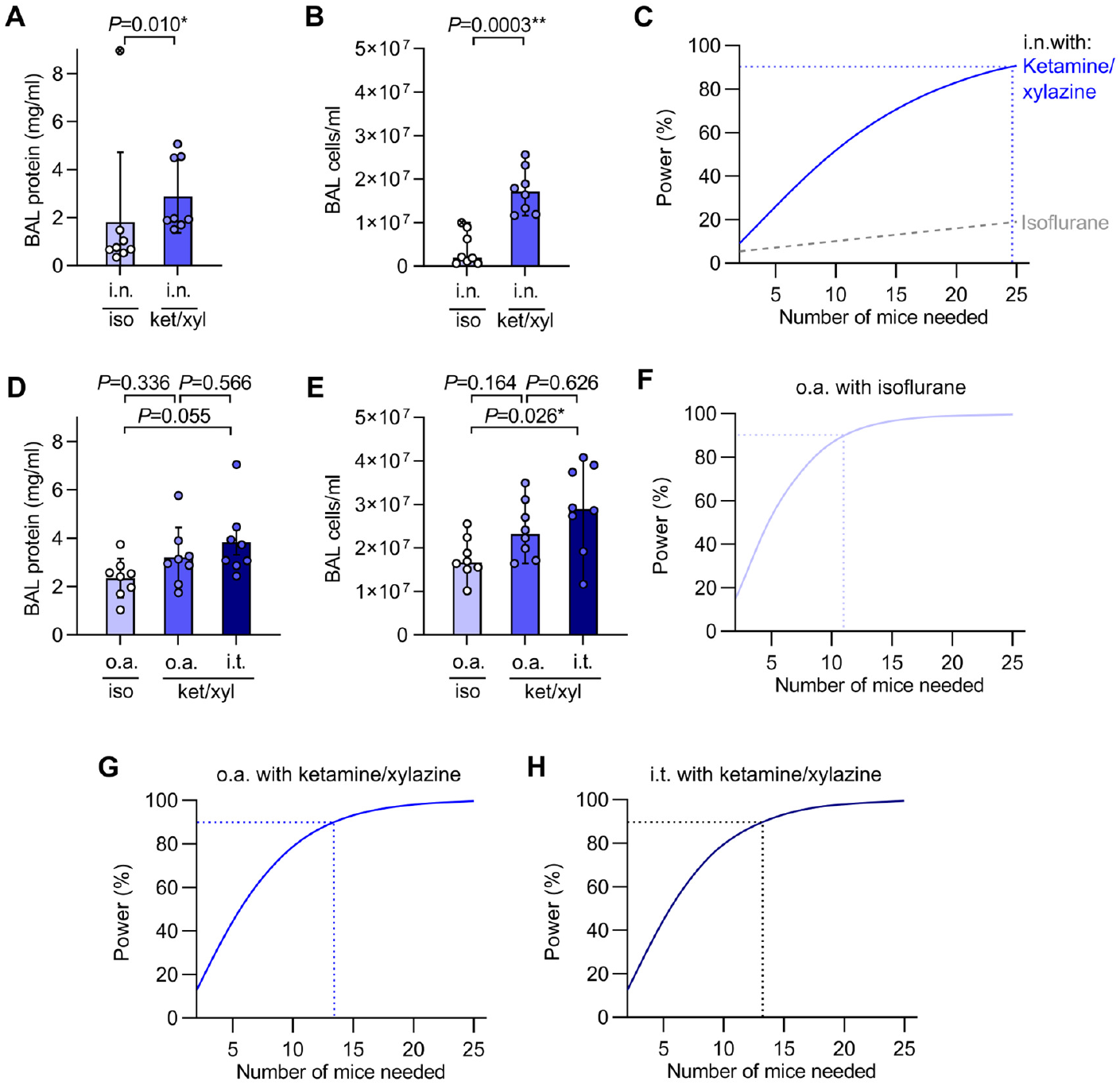
Anesthesia and administration approaches affect experimental power in a model of bacterial pneumonia. A. Mice were randomized to receive either isoflurane (iso) or ketamine/xylazine (ket/xyl) anesthesia for intranasal dosing with 1×10^6^ CFU of *Pseudomonas aeruginosa* (PAO1). BAL supernatant protein concentration was measured at 24 hours post infection. B. Cell counts from BAL fluid at 24 hours post infection. C. Output of power analysis using BAL protein data in Figure 5A to estimate number of mice needed per group to detect a 50% change in BAL protein concentration using an unpaired two-tailed t-test. E. Mice were randomized to receive 1×10^6^ CFU of PAO1 either via oropharyngeal aspiration (o.a.) with isoflurane anesthesia, o.a. with ketamine/xylazine anesthesia, or with intratracheal (i.t.) dosing under ketamine/xylazine anesthesia. BAL supernatant protein concentration was measured at 24 hours post infection. F. Cell counts from BAL fluid at 24 hours post infection. **F-H**. Output of power analyses using BAL protein data in Figure 5D to estimate number of mice needed per group to detect a 50% change in BAL protein concentration using an unpaired two-tailed t-test. Means ± standard deviation except for **B** and **E** which show medians ± 95% confidence intervals. Data from **B** and **E** were log_10_-transformed prior to analysis. *P*-values are from: **A**: Mann-Whitney U test; **B**: unpaired two-tailed t-test; **D**,**E**: ordinary one-way ANOVA with Tukey’s multiple comparisons test, n=8.

Relative to data with i.n. infection, delivery of the same bacterial inoculum via the o.a. or i.t. routes yielded greater increases in BAL protein and cell counts with lower variance (**Figure 5D, E**), giving increased experimental power (**Figure 5F-H**).

These results show how the improved pulmonary dosing efficiency achieved with the o.a. and i.t. methods relative to i.n. dosing can reduce the number of mice needed for experiments.

## Discussion

In this study, we found that anesthetic approach and administration route can affect the efficiency of fluid dosing to lungs of mice with impacts on experimental power in models of viral and bacterial pneumonia.

We conclude from our results that ketamine/xylazine anesthesia is preferable where consistent i.n. dosing to lungs of B6 mice is needed. As o.a. dosing is simple, can be achieved under isoflurane anesthesia and results in consistent pulmonary delivery, this route may be superior to i.n. dosing in some settings. Where precise dosing to lungs is critical, i.t. dosing might be useful as the i.t. administration approach that we describe resulted in high pulmonary dosing efficiency. Limitations of our i.t. dosing approach include the requirement for intubation and need for use of a nose cone or additional ketamine sedation for use with isoflurane anesthesia.

A likely consequence of poor delivery to lungs using i.n. dosing under isoflurane anesthesia is that inoculating doses containing greater quantities of virus will be used to produce infections that consistently result in robust lung inflammation, exposing extrapulmonary tissues to higher quantities of virus. This is not desirable, as exposure of the nasal sinuses to high quantities of viral particles could cause serious adverse effects, as recently demonstrated in a study showing lethal SARS-CoV-2 neuroinvasion when K18-hACE2 mice were infected using intranasal dosing but not when mice were infected using aerosolization to more gradually initiate an infection with smaller droplets while producing a similar pulmonary viral load (25). Our results in the bacterial lung infection model indicate that o.a. dosing might also be useful for improved modelling of viral pneumonia, and we speculate that retention and growth of bacteria in the nasal sinuses may also have contributed to the severe lung inflammation that developed in one of our mice given i.n. *P. aeruginosa* under isoflurane anesthesia.

Injectable ketamine/xylazine anesthesia may not always be preferable as recovery time can be longer than with isoflurane with potential effects on immune responses, and use of needles is discouraged where possible due to safety risks. In our study, we found that it was feasible to give mice i.p. ketamine/xylazine injections in a biosafety cabinet separate from that used for handling virus to minimize infection risk to handlers. Limiting dose reflux using ketamine/xylazine anesthesia might also reduce risk of aerosolization and surface contamination by inoculum, and animal suffering due to extrapulmonary pathology resulting from pathogen replication in the nasal sinuses (25).

Our study provides quantification of pulmonary dosing efficiency comparing i.n., o.a. and i.t. methods using anesthetics and mouse strains in current widespread usage. The percentage lung delivery values that we measured are largely consistent with those from previous studies which used a range of mouse strains, anesthetic approaches and methods for measuring dose distribution (1, 4, 14, 15). Our conclusion that anesthetic type can alter lung deposition of i.n. doses differs from that of a previous study which found no effect of different anesthetics (isoflurane, halothane, Avertin) during i.n. dosing on pulmonary dosing efficiency in BALB/c mice, although ketamine/xylazine was not examined in this previous report (1). Curiously, another study using BALB/c mice found increased bacterial content of lungs after intranasal dosing with *Francisella tularensis* under isoflurane compared to ketamine/xylazine anesthesia (26). B6 mice have wider bronchi than BALB/c mice (27), and *F. tularensis* given i.n. rapidly replicates outside of the lungs (28), so outcome of dosing may vary depending on mouse strain and pathogen biology.

In summary, we recommend the use of ketamine/xylazine anesthesia over isoflurane anesthesia for i.n. dosing into lungs of B6-background mice. Where needed, pulmonary dosing efficiency can be increased using the o.a. route, and further still using i.t. dosing. Anesthetic approach and administration method are factors that can alter outcomes of studies involving dosing to lungs, affecting the number of mice required for experiments. To increase reproducibility and decrease animal usage, these factors should be considered during experimental design and clearly reported in publications.

## Grants

We acknowledge support from:

Mark Looney: NIH R35HL161241 and R01AI160167.

Catharina Conrad: International Anesthesia Research Society Mentored Research Award.

S. Farshid Moussavi-Harami: NIH T32 Research Training in Pediatric Critical Care Medicine 2T32HD049303-16.

Simon Cleary: American Society of Transplantation Research Network/CSL Behring Basic Research Grant; Associated for the Advancement of Blood and Biotherapies Postdoctoral Grant; Sandler Program for Breakthrough Biomedical Research Postdoctoral Independence Grant.

## Author contributions

Data acquisition: Yurim Seo, Longhui Qiu, Simon Cleary. Conceptualization: Simon Cleary.

Methodology: Simon Cleary, Mélia Magnen, Catharina Conrad, S. Farshid Moussavi-Harami. Statistical analysis: Simon Cleary.

Resources: Mélia Magnen, Mark Looney, Simon Cleary. Writing – original draft: Yurim Seo, Simon Cleary.

Writing – review and editing: Yurim Seo, Mélia Magnen, Mark Looney, Simon Cleary Supervision: Simon Cleary, Mark Looney.

Funding acquisition: Mark Looney.

